# Human and chimpanzee-similar primates have distinct language gene polymorphism patterns

**DOI:** 10.1101/2023.07.20.549957

**Authors:** Wei Xia, Zhizhou Zhang

## Abstract

The difference in language gene polymorphism pattern (LGPP) between human and other primates may help to provide novel useful knowledge for language learning. One of important findings from many years’ worldwide research is that the primates like chimpanzee cannot easily recognize language grammars (even words). In this study, 189 SNPs (Single Nucleotide Polymorphism) in 13 language genes were scanned in 29 whole genomes from different human and primates populations. The 19 distinct SNPs in primates genomes were pointed out in several language genes including TPK1 that correlates with human’s syntactic and lexical ability. PCA analysis found that LGPPs for primates were highly aggregated together but they are distant from human’s LGPPs; representative human samples displayed high dispersion levels from each other in the context of LGPP. The above results may highlight a possibility that the LGPP should have more intermediate forms between human and chimpanzee-like primates.

## 1 Introduction

Chimpanzee’s learning ability, including language ability, has been investigated for many years [1-6], one of the aims of which is to understand why and how human performs much better than Primates. Especially, which differences in brain structures or (language) genes may contribute to the learning performance levels. Language ability has been a significant issue to investigate and compare among human individuals, chimpanzees (*Pan troglodytes*), Bonobos (*Pan paniscus*) and other primates. Language ability can be tested from listening, speaking to reading and writing. Apparently, listening itself may be not a problem for a chimpanzee, but speaking, reading and writing are too far away from chimpanzees’ capacity, though some other animals already possess ability for word regularities (affixation) [7-9]. It seems that this big difference derives only from the 1-2% difference in the genome sequences of human and chimpanzee. Communication between chimpanzees/bonobos in the wild takes the form of gestures, facial expressions, and a plenty of vocalization types, including grunts,roars, hoots, and screams. However, both kinds of Primates cannot orally speak like human even after many years’ education. The less dependence on tools, likely the less need to develop languages, because description of tool activities cannot be well fulfilled simply by non-oral expressions.

Language is a structured system that consists of grammar and vocabulary, by which humans convey meaning in the forms of spoken, written or signs of language. There are over 7000 different human languages with significant variations in cultural and historical diversity. Meanwhile, language has its own biological root. By now over a dozen of human language genes have been preliminarily characterized (Table 1). Though the human version of the gene FOXP2 harbours changes not found in chimpanzees or other primates, it is not reasonable to explain key language puzzles by a single mutation in modern humans [10,11]. But thousands of such mutations in language genes may help to find some patterns pointing to important issues, such as the cause of leaning ability difference. As the first step to this direction, this study employed 189 SNPs from 13 language genes in 29 whole genomes and found some SNP points that can significantly distinguish between human and primates.

**Table 1:** Language genes employed in this study.

| Name | Compromised language ability when mutated | Other functional information | References |
| --- | --- | --- | --- |
| ATP2C2 | Memory | Specific language impairment and some oral communication disorders | 21 |
| CMIP | Reading | Memory, speech and communication disorders | 22,23,21 |
| CNTNAP2 | Early language development | Implicated in multiple neurodevelopmental disorders, including schizophrenia, epilepsy, autism and intellectual disability | 22,24,25 |
| DCDC2 | Reading, dyslexia | Diseases associated with DCDC2 include deafness, autosomal recessive 66 and sclerosing cholangitis | 23,26,27 |
| FLNC | Reading, language | Involved in reorganizing the actin cytoskeleton in response to signaling events (such as vocalization) | 28 |
| FOXP1 | Expressive language | Diseases associated with FOXP1 include intellectual disability-severe speech delay-mild dysmorphism syndrome and intellectual developmental disorder with language impairment and with or without autistic features | 29 |
| FOXP2 | Speech |  | 30 |
| KIAA0319 | Reading, dyslexia | Involved in neuronal migration during development of the cerebral neocortex | 22,31,23,32 |
| NFXL1 | Speech | Specific language impairment | 33 |
| ROBO1 | Phonological buffer | axonal navigation at the ventral midline of the neural tube and projection of axons to different regions during neuronal development | 34,35 |
| ROBO2 | Expressive vocabulary | Nervous system development | 36 |
| TM4SF20 | Language delay | Neurobehavioral phenotype with abnormal motor coordination; communication disorder | 37 |
| TPK1 | Syntactic and lexical ability | Childhood encephalopathy | 38,39 |

**Table 2:** Tested 189 SNPs of thirteen language genes.

|  |  |  |  |  |
| --- | --- | --- | --- | --- |
| Language gene/SNP | Language gene/SNP | Language gene/SNP | Language gene/SNP | Language gene/SNP |
| ATP2C2 rs13334642 | CNTNAP2 rs987456 | FOXP1-rs1733518 | KIAA0319 rs7770041 | ROBO2 rs5788280 |
| ATP2C2 rs16973859 | DCDC2 rs190254728 | FOXP1-rs17803583 | NFXL1 rs1036681 | ROBO2 rs78817248 |
| ATP2C2 rs2435172 | DCDC2 rs2274305 | FOXP1-rs200643313 | NFXL1 rs12651301 | ROBO2-rs12171318 |
| ATP2C2 rs247818 | DCDC2 rs33914824 | FOXP1-rs2044341412 | NFXL1 rs13152765 | ROBO2-rs1372422 |
| ATP2C2 rs247885 | DCDC2 rs33943110 | FOXP1-rs2048059 | NFXL1 rs1371730 | ROBO2-rs1372427 |
| ATP2C2 rs4782948 | DCDC2 rs34584835 | FOXP1-rs722261 | NFXL1 rs1440228 | ROBO2-rs1503125 |
| ATP2C2 rs4782970 | DCDC2 rs35029429 | FOXP2 rs10227893 | NFXL1 rs147017712 | ROBO2-rs17203 |
| ATP2C2 rs62050917 | DCDC2 rs3789219 | FOXP2 rs10244649 | NFXL1 rs1545200 | ROBO2-rs264546 |
| ATP2C2 rs62640931 | DCDC2 rs3846827 | FOXP2 rs1058335 | NFXL1 rs1812964 | ROBO2-rs699456 |
| ATP2C2 rs62640932 | DCDC2 rs9460973 | FOXP2 rs12705977 | NFXL1 rs1822029 | ROBO2-rs873596 |
| ATP2C2 rs62640935 | DCDC2 rs9467075 | FOXP2 rs144807019 | NFXL1 rs1822030 | TM4SF20 rs13415654 |
| ATP2C2 rs74038217 | FLNC rs117864464 | FOXP2 rs182138317 | NFXL1 rs1964425 | TM4SF20 rs137891000 |
| ATP2C2 rs78371901 | FLNC rs2249128 | FOXP2 rs61732741 | NFXL1 rs34323060 | TM4SF20 rs4408717 |
| CMIP rs114894868 | FLNC rs2291558 | FOXP2 rs61753357 | NFXL1 rs920462 | TM4SF20 rs4428010 |
| CMIP rs1187121850 | FLNC rs2291560 | FOXP2 rs61758964 | NFXL1 rs978094 | TM4SF20 rs4438464 |
| CMIP rs16955675 | FLNC rs2291561 | FOXP2 rs62640396 | ROBO1 rs34841026 | TM4SF20 rs44675173 |
| CMIP rs183075361 | FLNC rs2291562 | FOXP2 rs73210755 | ROBO1 rs35456279 | TM4SF20 rs4673192 |
| CMIP rs183876152 | FLNC rs2291563 | FOXP2-rs531957198 | ROBO1 rs6795556 | TM4SF20 rs4675172 |
| CMIP rs201316817 | FLNC rs2291565 | FOXP2-rs718378 | ROBO1 rs77350918 | TM4SF20 rs6724955 |
| CMIP rs2288011 | FLNC rs2291566 | FOXP2-rs724419 | ROBO1-rs1378638 | TM4SF20 rs80305648 |
| CMIP rs34119643 | FLNC rs2291568 | FOXP2-rs747126499 | ROBO1-rs162423 | TM4SF20-rs10168278 |
| CMIP rs35429777 | FLNC rs2291569 | FOXP2-rs773664240 | ROBO1-rs331168 | TM4SF20-rs4675173 |
| CMIP rs57603843 | FLNC rs35281128 | FOXP2-rs776920 | ROBO1-rs3923148 | TM4SF20-rs7568026 |
| CMIP rs60152409 | FLNC rs371111092 | FOXP2-rs814066 | ROBO1-rs4130219 | TM4SF20-rs7574414 |
| CMIP rs74031247 | FOXP1 rs1053797 | FOXP2-rs940468 | ROBO1-rs4130431 | TM4SF20-rs9678000 |
| CMIP rs79979027 | FOXP1 rs11914627 | FOXP2-rs956016 | ROBO1-rs716681 | TPK1 rs113536847 |
| CNTNAP2 rs1062071 | FOXP1 rs144080925 | KIAA0319 rs10946705 | ROBO1-rs80030397 | TPK1 rs12333969 |
| CNTNAP2 rs1062072 | FOXP1 rs147756430 | KIAA0319 rs114195393 | ROBO1-rs991787 | TPK1 rs17170295 |
| CNTNAP2 rs1468370 | FOXP1 rs1499893 | KIAA0319 rs115399701 | ROBO2 rs1031377 | TPK1 rs28380423 |
| CNTNAP2 rs1479837 | FOXP1 rs17008063 | KIAA0319 rs117692893 | ROBO2 rs10865561 | TPK1 rs67644764 |
| CNTNAP2 rs1637841 | FOXP1 rs17008224 | KIAA0319 rs138160539 | ROBO2 rs11127602 | TPK1 rs6953807 |
| CNTNAP2 rs1637842 | FOXP1 rs17008544 | KIAA0319 rs150584710 | ROBO2 rs1163748 | TPK1 rs77358162 |
| CNTNAP2 rs2373284 | FOXP1 rs75214049 | KIAA0319 rs699461 | ROBO2 rs1163749 | TPK1 rs79464600 |
| CNTNAP2 rs3194 | FOXP1 rs76145927 | KIAA0319 rs699462 | ROBO2 rs1163750 | TPK1-rs228582 |
| CNTNAP2 rs535454043 | FOXP1 rs7638391 | KIAA0319 rs699463 | ROBO2 rs144468527 | TPK1-rs38045 |
| CNTNAP2 rs61732853 | FOXP1 rs7639736 | KIAA0319 rs730860 | ROBO2 rs17525412 | TPK1-rs38046 |
| CNTNAP2 rs700308 | FOXP1-rs1288693 | KIAA0319 rs75674723 | ROBO2 rs3923744 | TPK1-rs41239 |
| CNTNAP2 rs700309 | FOXP1-rs1463951 | KIAA0319 rs75720688 | ROBO2 rs3923745 |  |

### 1.1 Language ability is originally a complicated procedure of muscle movements

Language ability is one of the core features of the human species. The evolution of language ability should also witness the evolution of the human brain’s intelligence and the vocal organs, taking millions to tens of millions of years. The earliest primates record is about 60 million years. In 2022, it was reported that *Sahelanthropus tchadensis* (the known earliest human-like ape or early-stage *Homo erectus*) was able to walk upright 7 million years ago [12], and other studies inferred that late-stage *Homo erectus* should already be able to use language fluently [13]. The communication between apes is mainly gesture language [14], which goes through a long process with the evolution of the brain (and gradually solidified various gesture language and body language into a brain signal), and is reflected in the way of sound. Sound is also produced by the muscle movement of specific organs of the body, which is the mechanical movement of the occurring organs. This can also explain why language control regions in the brain are highly overlapping with motor control regions [15].

Evolution after upright walking: due to eating cooked food and genetic mutations, the brain organs have the ability to process complex signals; All of the organs are connected to the nervous system. The organs of the eyes, ears and mouth gradually produce a preliminary language; The movement of both hands also greatly promotes the generation of language; Language processes are actually similar to other limb movements, though language is the movement of the organs of the eyes, ears and mouth; The evolution process of language gene polymorphism is the process of the gradual improvement of language ability. The eyes, ears and mouths share the same neural control circuit as bimanual movements [15], because, in the course of evolution, the eyes, ears and mouths themselves are one of the most moving organs, or the most primitive motor organs. They received the most neural connections during evolution, laying the foundation for functional complexity. Meanwhile, technology for making and using tools in *Homo erectus* has also further evolved, and these techniques cannot be delivered precisely without the aid of language. Evolution may make the brain suddenly have a complex-enough computing power [16], opening an evolutionary opportunity for coordination between audiovisual stimulation signals and muscle movements, including opportunities to improve language levels.

### 1.2 Concepts of language gene and language gene polymorphism pattern (LGPP)

Genes directly related to language ability are called language genes. The direct correlation here means that if a gene is deleted or mutated or significantly altered in quantitative genetic traits, all or part of the language function is lost or weakened. At present, there are about 19 human language genes [13] [17] [18], with a large number of single nucleotide polymorphism (SNP) or mutation sites of single base sequence (SNV) on the sequence of each language gene. A total of 19 language genes can be several million SNP/SNV loci that can be used to describe the patterns of language gene polymorphism and their evolution dynamics in different ancient human samples.

### 1.3 LGPPs of human species and the primates

Research on advanced primates has been conducted for nearly a century. One of the main questions to clarify in such studies is understanding how higher primates such as chimpanzees evolved into humans. The differences between humans and chimpanzees, especially abilities in language and cognition are of primary concern. A large number of studies can be seen in some reviews [19,20], but few studies have been observed between the two from the perspective of language gene polymorphism patterns. In this study, 189 SNPs (single nuelotide polymorphism) in 13 language genes were scanned in 29 whole genomes from different human and primates populations.

## 2 Methods

### 2.1 Language genes and their SNPs

Table 1 listed 13 language genes (as a preliminary observation, only 13 language genes were employed at the time the manuscript was written), and a total 189 SNPs from these 13 genes were selected for this study. Language gene SNP data were all semi-randomly selected (Figure 1) for each gene in the dbSNP database: https://www.ncbi.nlm.nih.gov/snp/; Some SNPs have limited information on clinical effects as shown in dbSNP and GeneCards databases.

**Figure 1:**
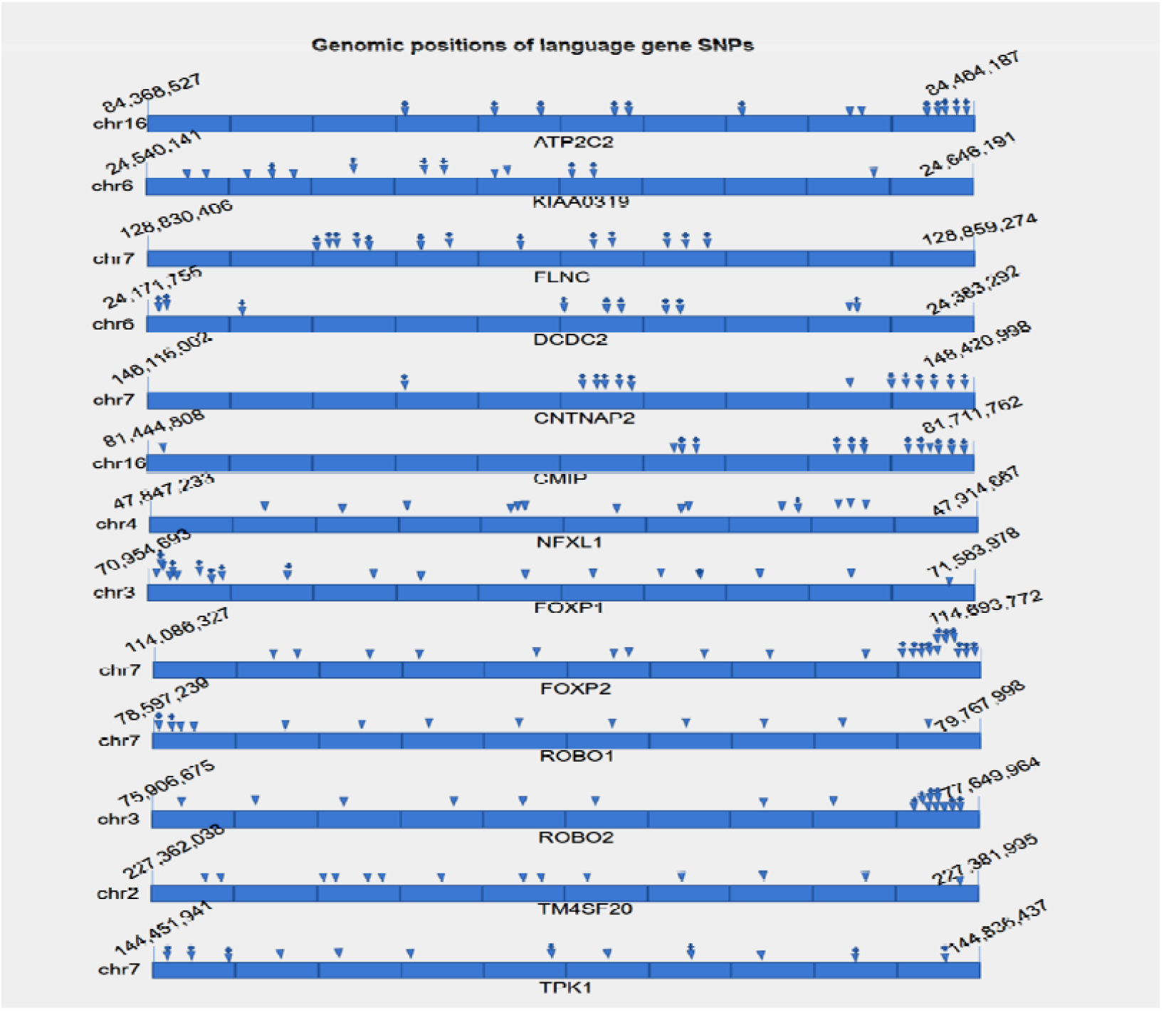
Semi-randomly selected SNP sites and their approximate positions in 13 language genes. The + sign above a triangle means the SNP site had a known clinical phenotype as indicated in dbSNP and GeneCards databases.

### 2.2 Genome sequences

All genome sequences (Table 3) were downloaded from ENA database (https://www.ebi.ac.uk/ena/browser/) in the fastq format. In all 29 genomes, the sizes mainly range from 41G to 200G. There are 14 representative human samples in which 10 were ancient samples. Four from Asia, one from Africa, 8 from Europe and one from South America. In the 15 primate samples, two from China, four from Indonesia and the left eight from Africa.

**Table 3:** The 29 whole genomes employed in this study.

| No. | abbr | Country | Region | Age (BP) | Detail | Genome file size (G) | References |
| --- | --- | --- | --- | --- | --- | --- | --- |
| 1 | pa6 | Pakistan | SouthAsia |  | Pakistan Brahui | 113 | PRJEB9586 |
| 2 | c19 | China (a) | EastAsia | 5304-5056 | China(WGM35) | 83 | PRJEB36297 |
| 3 | c9 | China (a) | EastAsia | 6175-5937 | XW-M1R18 | 117 | PRJEB36297 |
| 4 | dvi | China/Russia (a) | EastAsia | 8000 | DevilsGate | 41 | PRJEB14817 |
| 5 | ke2 | Kenya | Africa |  | Kenya Luhya-2 | 84 | PRJEB9586 |
| 6 | nd4 | Russia (a) | Europe | 50300 | Neanderthal Altai | 158 | PRJEB1265 |
| 7 | nd2 | Spain (a) | Europe | 60,000-120,000 | Neanderthal ForbesQuarry | 143 | PRJEB31410 |
| 8 | nd1 | Russia (a) | Europe | 50000 | Neanderthal-MIX1 | 64 | PRJEB29475 |
| 9 | sp1 | Spain | Europe |  | SPAIN-1 | 200 | PRJNA42557 |
| 10 | fr4 | France (a) | Europe | 4000 | France4,000 | 167 | PRJEB9586 |
| 11 | cz1 | Czech (a) | Europe | 45000 | Czechia ancient | 112 | PRJEB39040 |
| 12 | de2 | Russia (a) | Europe | 100000 | Denisova2 | 109 | PRJEB20653 |
| 13 | dep | Russia (a) | Europe | 74000-82000 | DenisovaPha | 95 | PRJEB3092 |
| 14 | pe1 | Peru | SouthAm |  | PERU ERR042535-MIX1 | 67 | PRJEB31736 |
| 15 | pp1 | Congo | Africa |  | Salonga Pan Paniscus | 63 | SRR741770<br>SRR741768<br>SRR741785 |
| 16 | pp2 | Congo | Africa |  | Pan Paniscus Kosana | 74 | PRJNA189439 |
| 17 | pp3 | Congo | Africa |  | Pan Paniscus Catherine | 105 | PRJNA189439 |
| 18 | rr1 | China | EastAsia |  | Rhinopithecus roxellanaRR0-RR5 + RR15-RR18 | 96 | PRJNA283338 |
| 19 | rr2 | China | EastAsia |  | Rhinopithecus roxellanaRR12-RR14 | 91 | PRJNA283338 |
| 20 | go1 | Congo | Africa |  | Gorilla-AZIZI | 89 | PRJNA189439 |
| 21 | go2 | Congo | Africa |  | Gorilla KAISI +SUZIE | 95 | PRJNA189439 |
| 22 | go3 | Congo | Africa |  | Gorilla Katie_B650 | 95 | PRJNA189439 |
| 23 | pg1 | Indonesia | SouthAsia |  | Pongo abelii Kiki | 87 | PRJNA189439 |
| 24 | pg2 | Indonesia | SouthAsia |  | Pongo abelii Elsi | 87 | PRJNA189439 |
| 25 | pg3 | Indonesia | SouthAsia |  | Pongo pygmaeus Sari | 102 | PRJNA189439 |
| 26 | pg4 | Indonesia | SouthAsia |  | Pongo tapanuliensis PA_B019 | 117 | PRJEB19688 |
| 27 | pt2 | Congo | Africa |  | Pan troglodytes Clint | 84 | PRJNA189439 |
| 28 | pt3 | Congo | Africa |  | Pan troglodytes Akwaya-Jean | 91 | PRJNA189439 |
| 29 | pt4 | Congo | Africa |  | Pan troglodytes Julie_LWC21 | 102 | PRJNA189439 |

Note: (a): ancient sample; SouthAm: South America;

### 2.3 SNP information extraction from genome sequences

This study was actually to investigate in what extent the primates and ancient human samples still hold the same SNP sequences as modern human, since the query sequences of SNPs were all downloaded from present dbSNP database that harbors mainly modern human SNPs. The authors used 010 Editor software to extract all 189 SNP information from each genome. For each SNP, about 20 left-flanking or right-flanking query nucleotides were searched in the genome in both DNA strands. Perfectly matched sequences were recorded using the SNP base A, T, G, C or combinations with them. For example, some sample may have both bases in a specific SNP sites, and thus recorded as AC, GC, etc. For those target sequences not perfectly matched around the SNP site, if the imperfect match occurs at the left flanking region, the SNP was recorded like A-, T-, C- or G-; if the imperfection occurs at the right flanking region, the SNP was recorded like A+, T+, C+ or G+ as in Table 5. Both-sides imperfection may occur such as A-G+, T-C- and A+C+. Imperfection counts only in the 4 bases left or right around the SNP site when at least one base in the four was mismatched (while the other bases in the query sequence were all matched). Any other cases of imperfection will be recorded as zero. The collected SNP data and the digitalized version can be requested from the authors.

### 2.4 PCA analysis with R codes

Principal Component Analysi(s PCA)was performed using R packages FactoMineR, factoextra and ggplot2.The main R codes are listed as follow. SNP data had to be digitalized before PCA performance. All SNP alleles were written in the sequence of A, T, G and C. For example, A, T, GC (not CG), TC (not CT) and ATC. For A,T,G and C, 999000000000, 999000000, 999000, and 999 were assigned, respectively. For two-letter SNP cases, such as AT, AC and GC, 999999000000, 999000000999 and 999999 were used, respectively. For those with left-flanking im-perfection, for example, A-,G- and C-, 997000000000, 997000 and 997 were assigned, respectively; for those with right-flanking im-perfection or both-sides im-perfection, such as T+, G+, A-C+, A+GC- and A+C-, 998000000, 998000, 997000000998, 998000999997 and 998000000997 were assigned, respectively.

~~~
> library(FactoMineR)
> library(factoextra)
> library(ggplot2)
> country <- read.delim(‘C:/RBook/20230315fastqSNPdata.txt’, row.names = 1, sep = ‘\t’)
> country <- t(country)
> country.pca <- PCA(country, ncp = 2, scale.unit = TRUE, graph = FALSE)
> plot(country.pca)
> pca_sample <- data.frame(country.pca$ind$coord[, 1:2])
> head(pca_sample)
> pca_eig1 <- round(country.pca$eig[1,2], 2)
> pca_eig2 <- round(country.pca$eig[2,2],2)
> pca_eig1
> pca_eig2
> group <- read.delim(‘C:/RBook/group3.txt’, row.names = 1, sep = ‘\t’, check.names = FALSE)
> group <- group[rownames(pca_sample),]
> pca_sample <- cbind(pca_sample, group)
> pca_sample$samples <- rownames(pca_sample)
> head(pca_sample)
> library(ggrepel)
> ggplot (data = pca_sample, aes (x = Dim.1, y = Dim.2)) + geom_point (aes (color = group), size
= 3) + scale_color_manual (values = c(‘purple’, ‘red’, ‘green’, ‘blue’, ‘brown’, ‘pink’, ‘yellow’,
‘orange’, ‘grey’)) + theme (panel.grid = element_blank(), panel.background = element_rect (color
= ‘black’, fill = ‘transparent’), legend.key = element_rect (fill=‘transparent’)) + labs(x=paste(‘PCA1:’,
pca_eig1, ‘%’), y=paste(‘PCA2:’, pca_eig2, ‘%’), color = ‘‘) + geom_text_repel (aes (label = samples),
size = 3, show.legend = FALSE, box.padding = unit(0.25, ‘lines’))
~~~

## 3 Result and discussion

There is a very high density between primate samples (figure 2), and there is a significant gap from the primates to human samples, though human samples are very dispersed. This result implies that either there are other intermediate species between primates and modern human, for example, different types of Homo erectus, or that the key LGPPs arise only after the emergence of *Homo erectus*. By now, the authors haven’t been able to collect genome sequences from fossil Australopithecus, including different types of *Homo erectus*. Hopefully their LGPPs just lie in the gap between the primates and modern human. There are some literature suggesting that the key events in language evolution occurred only in the hominin lineage, after the divergence from panins [19].

**Figure 2:**
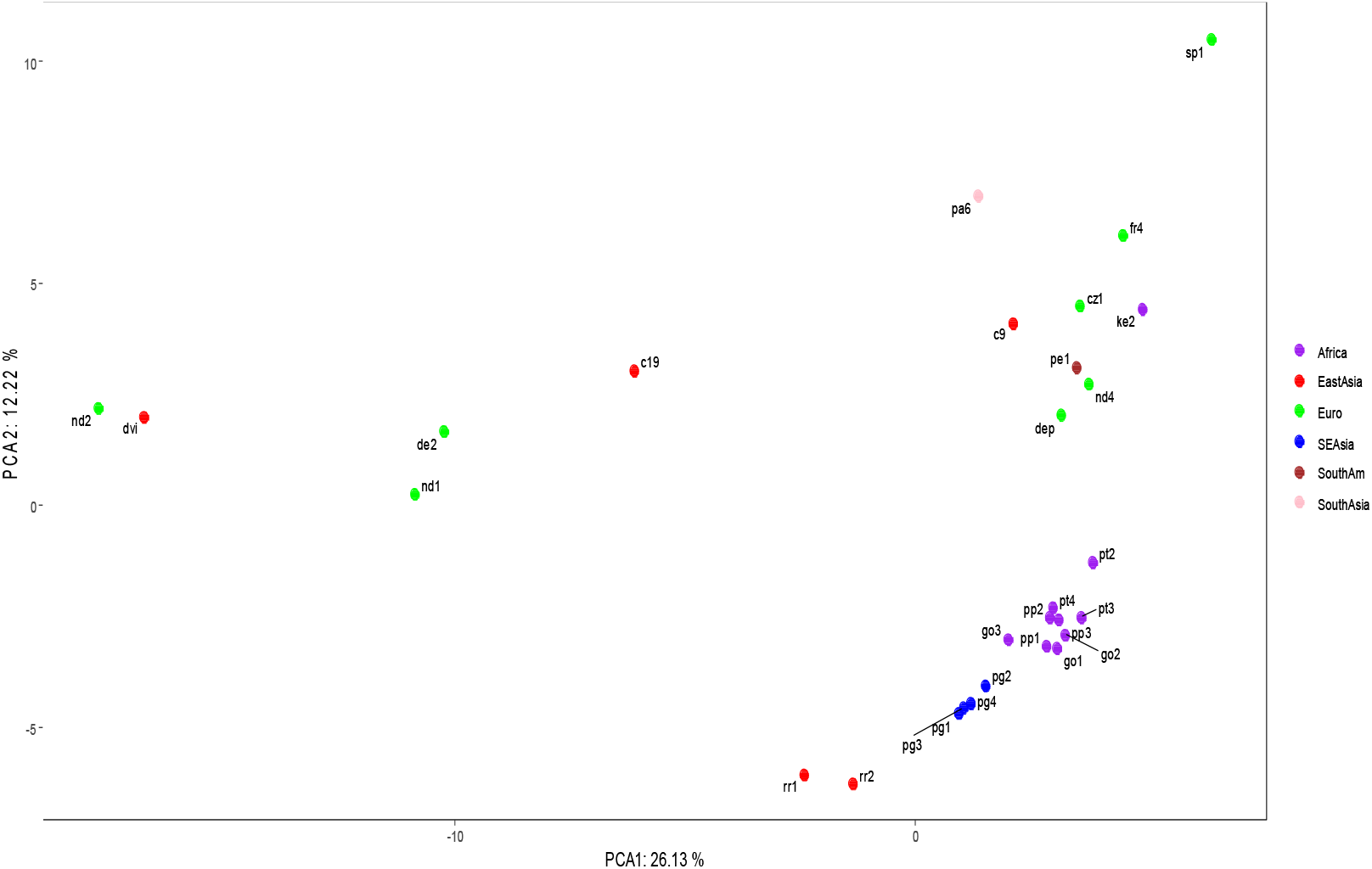
PCA demonstration for 29 samples in Table 3. Euro: Europe; SEAsia: South Asia; SouthAM: South America;

It is also possible that the language gene polymorphism pattern is not enough to distinguish humans from other animals, such as dolphins and parrots whose LGPPs are actually close to humans (data not shown). People probably have to rely on a mixed polymorphism pattern from both language genes and cognition genes. In theory, this hybrid model may be a better point to support the development of completely new language learning methods. This needs to be seen in much subsequent research.

Language evolution involves several important processes of Australopithecus-Man ape-Ape man-*Homo erectus*-*Homo sapiens*. Several key breakthroughs in language evolution were believed to be completed at some points in some above process. LGPP investigation may open a window for us to reach some breakthroughs. In Table 4, those SNPs with apparent difference between human and the primates were listed in nine language genes, and several SNP sites were significantly different between human samples and the primates, such as s1,s2,s3,s7,and s8 (also Table 5). The potential interaction network of the nine genes was described (Figure 3) with STRING tool in GeneCards database. Is that possible that human alleles and the primates alleles in the Figure 3 will lead to emergence of language ability’s great leap-forward? S7 and s8 refer to TPK1 gene, which has evidences to be associated with syntax and lexical retrieval [39], two important prerequisites for grammar function.

**Table 4:** Language genes employed in this study.

|  | pp3 | pp2 | pp1 | go3 | go2 | go1 | pg4 | pg3 | pg2 | pg1 | pt4 | pt3 | pt2 | pa6 | ke2 | SP1 | PE1 | FR4 | c19 | c9 | Dvi | De2 | Cz1 | Nd1 | Nd4 | Nd2 | DeP |
| --- | --- | --- | --- | --- | --- | --- | --- | --- | --- | --- | --- | --- | --- | --- | --- | --- | --- | --- | --- | --- | --- | --- | --- | --- | --- | --- | --- |
| s1 | A | A | A | A | A | A | A- | A | A | A | A | A | A | G | G | TG | G | T | 0 | 0 | 0 | 0 | G | 0 | G | 0 | G |
| s2 | C | C | C | C | C | C | C | C | C | C | C | C | C | TC | C | TC | CT | C | 0 | C | 0 | 0 | C | 0 | TC | 0 | T |
| s3 | G | G | G | G | G | G | G | G | G | G | G | G | G | C | GC | C | C | C | C | GC | 0 | 0 | GC | C | C | 0 | C |
| s4 | G | G | G | G+ | G+ | G+ | G | G | G | G | G | G | G | G+ | A | AG | A | A | G | AG | A | G | AG | A | A | 0 | G |
| s5 | G | G | G | A | A | A | G | G | G | G | G | G | G | G | G | G | G | AG | G | G | 0 | 0 | G | 0 | G | 0 | G |
| s6 | C | C | C | C | C | C | C | C | C | C | C | C | C | TC | T | TC | C | C | C | C | 0 | C | C | 0 | C | 0 | C |
| s7 | T | T | T | AT | T | T | T | T | T | T | T | T | T | T | G | G | G | G | T | 0 | 0 | 0 | G | G | G | 0 | G |
| s8 | T | T | T | T | T | T | T | T | T | T | T | T | T | TG | T | TG | G | G | G | G | T | G | G | T | G | 0 | G |
| s9 | G | G | G | G | G | G | G | G | G | G | G | G | G | AG | G | AG | G | G | A | 0 | 0 | G | G | G | G | 0 | G |
| s10 | C | C | C | C | C | C | C | C | C | C | C | C | C | A | AC | AC | A | AC | 0 | 0 | 0 | 0 | A | C | AC | 0 | C |
| s11 | G | G | G | G | G | G | G | G | G | G | G | G | G | TG | G | TG | G | TG | G | G | 0 | G | T | 0 | G | G | G |
| s12 | A | A | A | A | A | A | A | A | A | A | A | A | A | C | AC | AC | A | A | 0 | A | 0 | 0 | C | C | C | 0 | C |
| s13 | G | G | G | G | G | G | G | G | G | 0 | G | G | G | TG | G | TG | G | 0 | 0 | G | 0 | G | T | 0 | G | 0 | G |
| s14 | G | G | G | G | G | G | G | G | G | G | G | G | G | 0 | G | G | G | G | A | G | 0 | 0 | G | 0 | G | 0 | G |
| s15 | C | C | C | C | C | C | C | C | C | C | C | C | C | A+ | C | C | C | C | C | C | 0 | C | C | T | C | 0 | C |
| s16 | 0 | G | G | 0 | A | A | G | G | G | G | G | G | G | 0 | G | G | G | G | 0 | G | C | G | G | G | G | 0 | G |
| s17 | C | C | C | C | C+ | C | 0 | 0 | 0 | 0 | C | C | C | 0 | C | C | C | TC | C | C | C | 0 | C | C | C | 0 | C |
| s18 | A | A | A | G | G | G | 0 | 0 | 0 | A | A | A | A | 0 | G | G | G | G | G | G | 0 | 0 | G | G | G | 0 | G |
| s19 | A | A | A | AT | A | T | A- | A | 0 | A | A | A | A | A | AG | G | A | A | 0 | 0 | 0 | A | G | G | G | 0 | A |

Note: The yellow-colored parts refer to SNPs that are significantly different between Primates and human samples. Blue colors denote some Gorilla- or Pongo-specific information.

**Table 5:** Language genes employed in this study.

| SNP | SNP |
| --- | --- |
| s1† ROBO1 rs6795556*<sup>t2</sup> | s11 CNTNAP2 rs1468370*<sup>t1</sup> |
| s2† ROBO2 rs10865561*<sup>t3</sup> | s12 CNTNAP2 rs987456*<sup>t1t4</sup> |
| s3† TM4SF20 rs6724955<sup>t1</sup> | s13 CNTNAP2 rs700309*<sup>t1</sup> |
| s4 TM4SF20 rs4438464<sup>t1</sup> | s14 CMIP rs183876152*<sup>t2</sup> |
| s5 TPK1 rs113536847*<sup>t4</sup> | s15 CMIP rs114894868*<sup>t2</sup> |
| s6 TPK1 rs28380423*<sup>t1</sup> | s16 ATP2C2 rs78371901*<sup>t2</sup> |
| s7† TPK1 rs17170295*<sup>t1</sup> | s17 ATP2C2 rs62050917*<sup>t5</sup> |
| s8† TPK1 rs67644764<sup>t1t2</sup> | s18 ATP2C2 rs4782948*<sup>t2</sup> |
| s9 NFXL1 rs1822030<sup>t1</sup> | s19 KIAA0319 rs699461<sup>t1</sup> |
| s10 KIAA0319 rs699461<sup>t1</sup> |  |

\*benign clinical significance; t1: intron variant; t2: coding sequence variant; t3: downstream
transcript variant; t4: 3' UTR variant; t5: non coding transcript variant; †: significant difference
between human and Primates;

**Figure 3:**
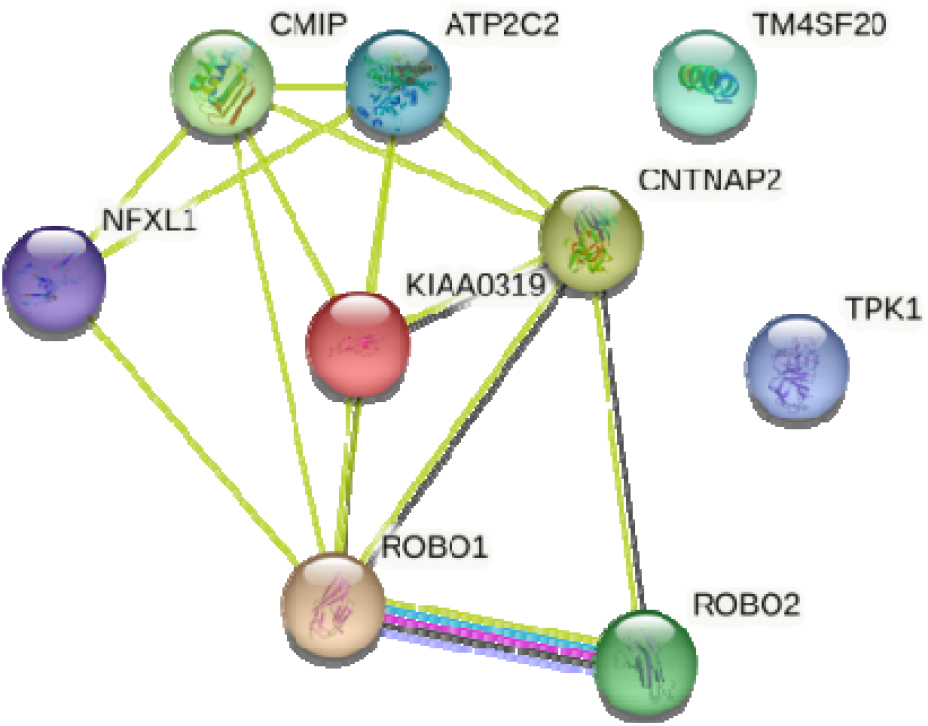
Potential molecular interactions among nine genes (as proteins) in Table 5. TPK1 and TM4SF20 were not linked in the potential network. Colors of the lines between two genes represent interaction types (see String database).

Since language ability was first just a muscle movement behavior, from this perspective, language must have evolved much earlier than human evolution. This is evident from other results (data not shown) of our preliminary study. We found that the language gene polymorphism pattern of parrots and dolphins is very close to that of some ancient humans and some modern humans, but the language gene polymorphism pattern of advanced primates such as chimpanzees is obviously or relatively far from human beings; From the ratio of brain weight to body weight, the human brain accounted for 2.1% of body weight, dolphins 1.17%, and chimpanzees only 0.7%. The fossils of parrots reached 55 million years [40]; the 50 million year ancestor of Dolphin, Pakicetus [41], was found in Pakistan, and the dolphin family emerged from the Miocene about 12 million years ago. But the earliest known upright Homo species was Chad Australopithecus only 7 million years ago. So these data indicate that some language gene polymorphism patterns appeared in the evolution much earlier than dolphins and other animals. If this is true, LGPP itself may be not enough to explain language ability difference between human and chimpanzees.

## 4 Conclusion

In this study, 189 semi-randomly selected SNPs (Single Nucleotide Polymorphism) in 13 language genes (SNP positions seen in figure 1) were scanned in 29 whole genomes from different human and primates populations. The 19 distinct SNPs in primates genomes were found in several language genes including TPK1 that correlates with human’s syntactic and lexical ability.

PCA result indicated that the language gene polymorphism pattern does demonstrate differences between primates and representative human samples, and the difference appears quite obvious. This difference suggests that there should be some intermediate states between primates and modern humans, presumably among the late-stage apes, *Homo erectus*, or early *Homo sapiens*. But these intermediate species samples are the most scarce, usually very precious fossil-bone, skeleton or tooth samples. Given that investigations on language gene polymorphism patterns have found that LGPP is more similar (than the primates) between dolphins and human samples, we may not be able to find human-specific LGPP in the future even if we have enough fossil *Homo erectus* genome sequences and the corresponding LGPPs. It is likely that the polymorphism patterns of language genes and cognitive genes will need to be mixed together for a broader investigation in the future study.

## ACKNOWLEDGMENTS

This study was supported by State Language Commission Research Grant (YB135-117) and National Research Center for Foreign Language Education Grant (ZGWYJYJJ10A042).

